# Codon Use and Aversion is Largely Phylogenetically Conserved Across the Tree of Life

**DOI:** 10.1101/649590

**Authors:** Justin B. Miller, Lauren M. McKinnon, Michael F. Whiting, Perry G. Ridge

## Abstract

Using parsimony, we analyzed codon usages across 12 337 species and 25 727 orthologous genes to rank specific genes and codons according to their phylogenetic signal. We examined each codon within each ortholog to determine the codon usage for each species. In total, 890 814 codons were parsimony informative. Next, we compared species that used a codon with species that did not use the codon. We assessed each codon’s congruence with species relationships provided in the Open Tree of Life (OTL) and determined the statistical probability of observing these results by random chance. We determined that 25 771 codons had no parallelisms or reversals when mapped to the OTL. Codon usages from orthologous genes spanning many species were 1 109x more likely to be congruent with species relationships in the OTL than would be expected by random chance. Using the OTL as a reference, we show that codon usage is phylogenetically conserved within orthologous genes in archaea, bacteria, plants, mammals, and other vertebrates. We also show how to use our provided framework to test different tree hypotheses by confirming the placement of turtles as sister taxa to archosaurs.

**Availability:** All scripts, a README, and necessary test files are freely available on GitHub at https://github.com/ridgelab/codon_congruence

## 1. Introduction

The genetic code is degenerate because 64 canonical codons encode 20 amino acids and the stop codon, meaning multiple synonymous codons encode the same amino acid (Crick, 1970; Crick, 1966, 1968; Crick et al., 1961). Codon usage bias refers to the unequal distribution of synonymous codons between species, genes, or locations within the same gene, and can be used to regulate gene expression (Quax et al., 2015), suggesting that codon choice, even when synonymous, has biological implications. Typically, more closely related species share more similar patterns of codon usage and codon aversion (i.e., when a species does not use a codon within an ortholog), and these patterns are phylogenetically conserved (Miller et al., 2017). However, similar to other genetic characters (Rokas and Carroll, 2008), parallelism is present in the usage or aversion to many codons, resulting in homoplasy. In codon data, homoplasy may occur by parallelism, convergence, or reversal, resulting in identical character states that were not directly inherited from the most recent common ancestor. The presence of homoplasy is the greatest challenge in phylogenetic estimation, and nearly all characters, whether morphological or molecular, display homoplasy of some form at some level (Sanderson and Hufford, 1996).

To limit the effects of contradictory signals from homoplasy, the commonly-used maximum likelihood statistical method for estimating phylogenies approximates rates of evolution (e.g., transition and transversion ratios, evolutionary clock, evolutionary distance of species, etc.) and tree topography (Felsenstein, 1981). The basis of maximum likelihood is the proportionality of the likelihood function to the multinomial probability of observing the data given the tree and model (Huelsenbeck and Crandall, 1997; Yang et al., 1994). Maximum likelihood also uses the statistical property of consistency, which shows that as the number of data points approaches infinity, the maximum likelihood estimators will converge on the same estimate (Wald, 1949). In contrast, parsimony does not use a model to recover phylogenies, which potentially limits the consistency of the method (Felsenstein, 1978); however, *ad hoc* hypotheses of homoplasy are also limited (Farris, 1983).

From a likelihood standpoint, a model of codon usage and codon aversion requires understanding how codon usages change through evolutionary time. Various codon usage models have previously been proposed. Divergent selective pressures at specific amino acid sites have also been identified with maximum likelihood (Bielawski and Yang, 2004). A popular tool called PAML 4 (Yang, 2007) allows users to compare different evolutionary models, estimate synonymous and nonsynonymous mutation rates, infer selection, and reconstruct ancestral genes by utilizing a suite of codon models including Markov-based estimates of codon substitution rates (Goldman and Yang, 1994) and models based on instantaneous substitution rates being proportional to the frequency of a specific nucleotide (Muse and Gaut, 1994). CodonPhyML (Gil et al., 2013) estimates phylogenetic branch support based on hundreds of codon models. Although significantly faster than other algorithms, CodonPhyML is capable of analyzing only a few hundred species because of the computational complexity of maximum likelihood. Because of the high computational cost of maximum likelihood-based approaches, domain partitioning at predefined locations in the data have been used to allow for analyzing the data using multiple codon usage models (Zoller et al., 2015).

These previous models based on codon usage bias provide a strong foundation for the use of codons in a phylogenetic framework. However, these analyses based on codon usage bias may be complicated by additional confounding factors. For example, distinct domains with different codon usages exist, including a ramp of slowly translated codons at the beginnings of gene sequences (Miller et al., 2019; Tuller et al., 2010). In our analysis, we analyze both codon usage and codon aversion to account for these factors.

Although several models of codon evolution have been implemented, their use in a maximum likelihood framework is impractical across the Tree of Life using whole genome sequencing. Therefore, we must first start with parsimony. From a parsimony perspective, a tree hypothesis is meant to minimize the number of similarities left unexplained (Farris, 2008). We aimed to determine the extent that codon usage and codon aversion within orthologous genes is congruent with species relationships as presented in the Open Tree of Life (OTL) (Hinchliff et al., 2015).

Within genes, codon usage regulates gene expression in various ways. Using an equal number of codons to the supply of cognate tRNA anticodons maintains optimal codon usage that increases translational efficiency (Sharp and Li, 1987). Using multiple instances of the same codon (identical codon pairing) or synonymous codons (co-tRNA codon pairing) within a ribosomal window also increases translational efficiency and speed (Cannarozzi et al., 2010). Furthermore, mRNA structural folding and differential protein production are affected by codon usage bias within a gene (Gingold et al., 2014; Pechmann and Frydman, 2013; Quax et al., 2015). Therefore, codon usage within a gene, although potentially not homologous at a given position, is homologous on a genic level and can influence mRNA folding, protein production, and tRNA translational efficiency.

Encoding the codon matrix based on codon usage anywhere in the gene was first proposed by Miller et al. (2017), and categorizes homology on a genic scale, instead of positional homology from a multiple sequence alignment. This method uses a binary representation of codon usage (i.e., if the codon is used in a given ortholog then it is encoded as a ‘1’ and if it is not used, then it is encoded as a ‘0’). This process determines if a species “chooses” to use a given codon within a gene. Their results suggest that codon aversion is a parsimony informative character state in tetrapods. Furthermore, they show that using only stop codons recovers a phylogenetic tree that most closely resembles currently proposed phylogenies. Although Miller et al. (2017) established a basis for codon aversion in a phylogenetic framework, their study was limited to 72 tetrapods, and they did not establish a method to use codon aversion to resolve controversial nodes. In the current study, we show that codon aversion is phylogenetically conserved by analyzing 12 337 species across all domains of life. We further apply conditional probability of codon usages to different tree topologies to assess the probability of each tree occurring by random chance. We apply this method to assess the placement of turtles on the phylogeny, lending support to a recent representation of these taxa by Shen et al. (2017).

## 2. Materials and Methods

### 2.1 Data Collection and Processing

We downloaded 12 337 reference genomes, one for each species, and their accompanying General Feature Format (GFF3) files from the National Center for Biotechnology Information (NCBI) (Coordinators, 2013; Pruitt et al., 2014; Pruitt et al., 2000; Tatusova et al., 2014) in September 2017 using their FTP site: ftp://ftp.ncbi.nlm.nih.gov/genomes/refseq/. A reference genome represents the consensus genome for a species based on the most complete genome assemblies (Pruitt et al., 2014). We extracted all coding sequence (CDS) data from the reference genomes, and we assigned each of the 12 337 species to the following groups: 362 archaea, 11 227 bacteria, 214 fungi, 147 invertebrates, 105 mammals, 120 other vertebrates, 87 plants, and 75 protozoa based on species annotations in NCBI. Since viruses are not included in the OTL, they were not included in our analysis. We recognize that several of these taxonomic groups do not represent monophyletic clades, but we opted to keep the groups outlined in the NCBI database to facilitate comparisons between studies that also use these annotations. We required that CDS regions be annotated with an NCBI gene name, which combines gene annotations from species-specific nomenclature committees (e.g., the HUGO Gene Nomenclature Committee (HGNC) (Gray et al., 2015)), to ensure that orthologous comparisons of codon usage were used. Although we do not perform any formal analysis to verify the orthologous relationships proposed by NCBI, NCBI standardizes various gene studies, NCBI staff curations, and the NCBI annotation pipeline with gene annotations in SWISS-PROT (UniProt Consortium, 2018) to facilitate ortholog comparisons between species.

Next, we filtered the CDS regions to remove any annotated exceptions (e.g., translational exceptions, unclassified transcription discrepancies, suspected errors, etc.), as defined in the “attributes” column of the GFF3 files that were downloaded from NCBI. We used the longest isoform of each gene when multiple isoforms were annotated in order to include all codons that are used in the gene. We also included partial gene sequences where the orthologous relationship was annotated in order to include as many orthologous gene comparisons as possible. Finally, we required each ortholog to be present in at least four species to ensure that the codon usage could be parsimony informative.

For each codon within each ortholog, we encoded its usage in a binary matrix (i.e., if the codon was used, it was given a ‘1’ and if it was not used, it was given a ‘0’). After all codons within all orthologs for all species were included in the binary matrix, we filtered out parsimony uninformative characters (i.e., when all species with an ortholog either used or did not use a given codon). For each remaining codon, we divided the species sampled into two partitions based on their character state for that codon: species that use a codon within an ortholog and species that do not use that codon within the same ortholog. This process is depicted in Figure 1.

**Figure 1:**
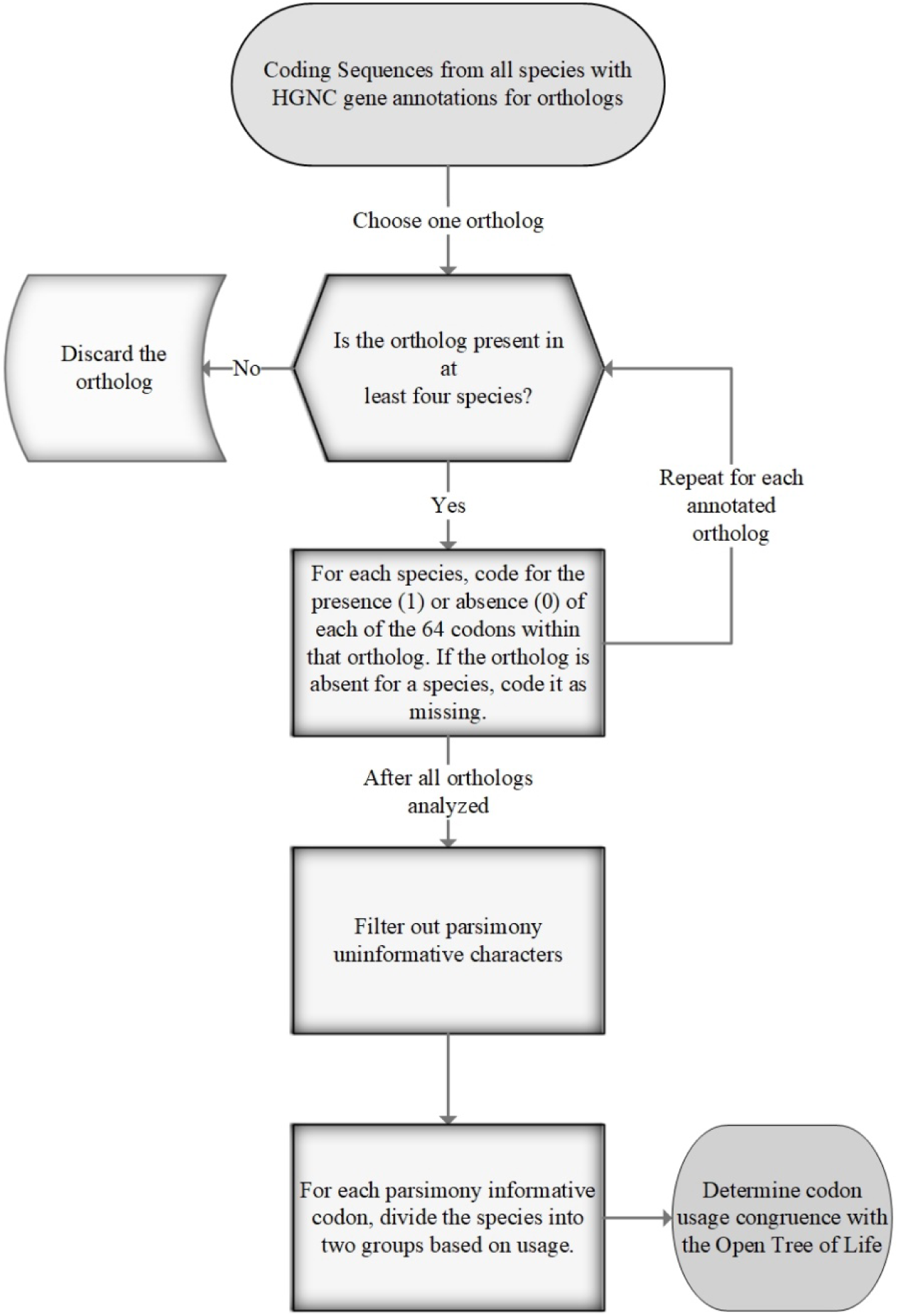
The process for encoding codon usage. Codon characters are encoded as either present (1) or absent (0) if they are used or not used in an ortholog, respectively.

After encoding the binary matrix for each codon within each ortholog, we evaluated each bipartition against the OTL to determine if parallelisms or reversals occurred. Parallelisms occur when the same codon independently arises in different lineages not due to a common ancestor. Reversals occur when a codon reverts back to an ancestral state (e.g., if a species uses a codon that its most resent ancestor did not use, but the codon was used by a more distant ancestor). For each orthologous gene and codon state in each bipartition, we report the number of gains, losses, unknown gain/loss at the root node, number of species in the smaller partition, total species with that ortholog, percent of species in the smaller partition, and total number of gain/loss divided by the number of species in the smaller partition (see Supplementary Tables 1-9). A codon was classified as separating species according to taxonomic groups reported in the OTL if the smaller group had at least two species and the total number of gain/loss and unknown gain/loss at the root node equaled one. This process was used because a singular gain/loss event would indicate that no reversals or parallelisms occurred for that codon character state and its state is unique to that lineage.

**Table 1:** The probability of codons mapping to the OTL tree topology due to random chance assuming no phylogenetic signal in codon usage. The first column shows the species divisions, with the first row being a combination of all species. The second column shows the number of taxonomic distributions. The third column shows the number of codon characters that completely follow species relationships shown in the OTL. The fourth column shows the t-statistic obtained from performing a chi-square test on the expected number of congruent characters versus the actual number of congruent characters, with respect to the OTL. The fifth column shows the p-value of the data, obtained from the t-statistic and the degrees of freedom from the number of different taxonomic distributions. The sixth column is the best t-statistic obtained from 100 random permutations of the species while maintaining the same tree structure. The seventh column is the p-value obtained from the highest t-statistic from 100 random permutations of the species while maintaining the same tree structure as the OTL. The eighth column shows the number of random permutations where the permutated p-value is ≤ the observed p-value.

| Taxonomic Group | Number of Different Taxonomic Distributions | Number of Codons with no Parallelisms or Reversals OTL | *t*-statistic | *p*-value | Best Random Permutation *t*-statistic | Best Random Permutation *p*-value | Number of Random Permutations with *p*-value is less than or equal to the observed |
| --- | --- | --- | --- | --- | --- | --- | --- |
| All | 62,416 | 25,771 | 4.12488x10<sup>46</sup> | 0 | 1.77991x10<sup>3</sup> | 1.0 | 0 |
| Archaea | 2,320 | 925 | 8.28913x10<sup>20</sup> | 0 | 2.93471x10<sup>3</sup> | 3.26479x10<sup>-17</sup> | 0 |
| Bacteria | 58,368 | 6,639 | 1.53961x10<sup>13</sup> | 0 | 1.93163x10<sup>3</sup> | 1.0 | 0 |
| Fungi | 16 | 2,019 | 2.55445x10<sup>2</sup> | 9.37392x10<sup>-46</sup> | 5.99657x10<sup>2</sup> | 4.19847x10<sup>-118</sup> | 16 |
| Invertebrates | 182 | 124 | 2.78440x10<sup>2</sup> | 4.34253x10<sup>-6</sup> | 6.54011x10<sup>2</sup> | 9.54888x10<sup>-55</sup> | 3 |
| Plants | 477 | 1,702 | 3.90146x10<sup>9</sup> | 0 | 1.39101x10<sup>3</sup> | 1.89724x10<sup>-90</sup> | 0 |
| Protozoa | 21 | 2,449 | 2.14626x10<sup>3</sup> | 0 | 6.80127x10<sup>2</sup> | 3.53259x10<sup>-131</sup> | 0 |
| Mammals | 2,162 | 10,029 | 1.32695x10<sup>22</sup> | 0 | 4.04197x10<sup>3</sup> | 3.34856x10<sup>-117</sup> | 0 |
| Other Vertebrates | 2,770 | 11,877 | 1.50814x10<sup>25</sup> | 0 | 1.16996x10<sup>3</sup> | 1.0 | 0 |

### 2.2 Statistical Validation

Because autapomorphies are not parsimony informative, we required that the smaller partition include at least two species before it was mapped to the OTL. We then determined where on the OTL species gain or lose the usage of each codon character. Initially mapping the codon usage from a single species to the OTL has a probability of 1.0 of mapping to a taxonomic group that is congruent with the OTL because autapomorphies provide no evidence of species relationships. If the remaining character-state distributions randomly separate the other species (i.e., the null hypothesis), then we can use conditional probabilities to calculate the probability that a monophyletic group of more than one species is obtained by random chance. In this case, the probability that another species from the same taxonomic group as the first species would be correctly added to the same taxonomic group as the first, or subsequent, species is given in equation (1).

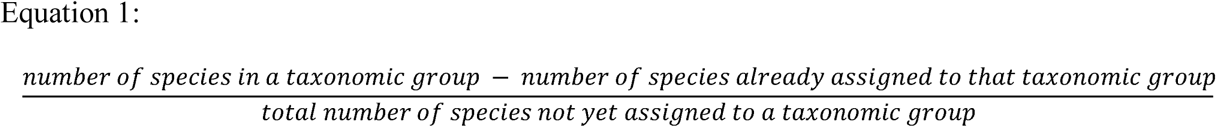

Using conditional probability, we calculate the probability of correctly assigning all members of a species partition to the taxonomic groups outlined in the OTL by random chance, using Equation (2). In Equation (2) *s*=the number of species in the smaller taxonomic group, and *t*=the total number of species sampled. We start with *s*-1 and *t*-1 to account for the initial species that was mapped to the phylogeny, which will always have a probability of 1.0 that it is correctly placed in a monophyletic group.

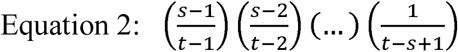

This equation simplifies to equation (3).

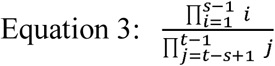

A taxonomic distribution is defined as the number of species in the sets separated by the codon character state (i.e., the number of species that use a given codon and the number of species that do not use a given codon within an ortholog), without regard to the OTL. Using equation (3), we calculated the expected number of significant character states for each taxonomic distribution (e.g., if five species were sampled, with two species in the smaller group, then, using equation (3), the probability that they were correctly divided by random chance is 0.25). We then multiplied the probability of that taxonomic distribution correctly agreeing with the OTL by the number of instances of that taxonomic distribution in our dataset (e.g., if 12 instances of five species dividing into groups of two and three occurred in our dataset, then the expected number of species partitions agreeing with the OTL taxonomic groups would be 0.25 * 12 = 3). We performed a chi square analysis for the taxonomic distributions using these expected values and the observed values from our analysis.

### 2.3 Random Permutations

The statistical validation gives a theoretical probability of obtaining these results by random chance. However, it does not take into account the degree of homoplasy in the dataset or the overall congruence within the homoplastic character states with respect to other trees than the OTL. For these reasons, random permutations are needed to ensure that the probability of character states mapping to random trees does not exceed the probability of the observed character states mapping to the OTL.

Random permutations were conducted 100 times for each taxonomic group. Each permutation maintained the tree structure as hypothesized in the OTL to not bias our results based on artificial tree structures. We then randomized the distribution of the species in each taxonomic group, creating 100 different species relationships using the same tree structure. Next, we conducted the same statistical validation on each of these trees with randomly distributed species, as outlined above. We calculated the number of permutations where the *p-value* of the mapped characters was less than or equal to the observed *p-value*. Where random permutations obtained a smaller *p-value* than the observed, we concluded that there was not support for codon usage as being more congruent with the OTL than expected by random chance.

### 2.4 Visualizing Homology on the Tree of Life

We inferred the reference phylogenies from the OTL for each pre-defined taxonomic group using tools available in the OTL documentation. We then mapped each character state to the inferred subtree from the OTL and determined how many gains, losses, parallelisms, and reversals occurred. The entire process of mapping character states from the original coding sequences to the phylogeny in Newick format is outlined in Figure 2. Visualizations of the phylogenies were created using FigTree (http://tree.bio.ed.ac.uk/software/figtree/).

**Figure 2:**
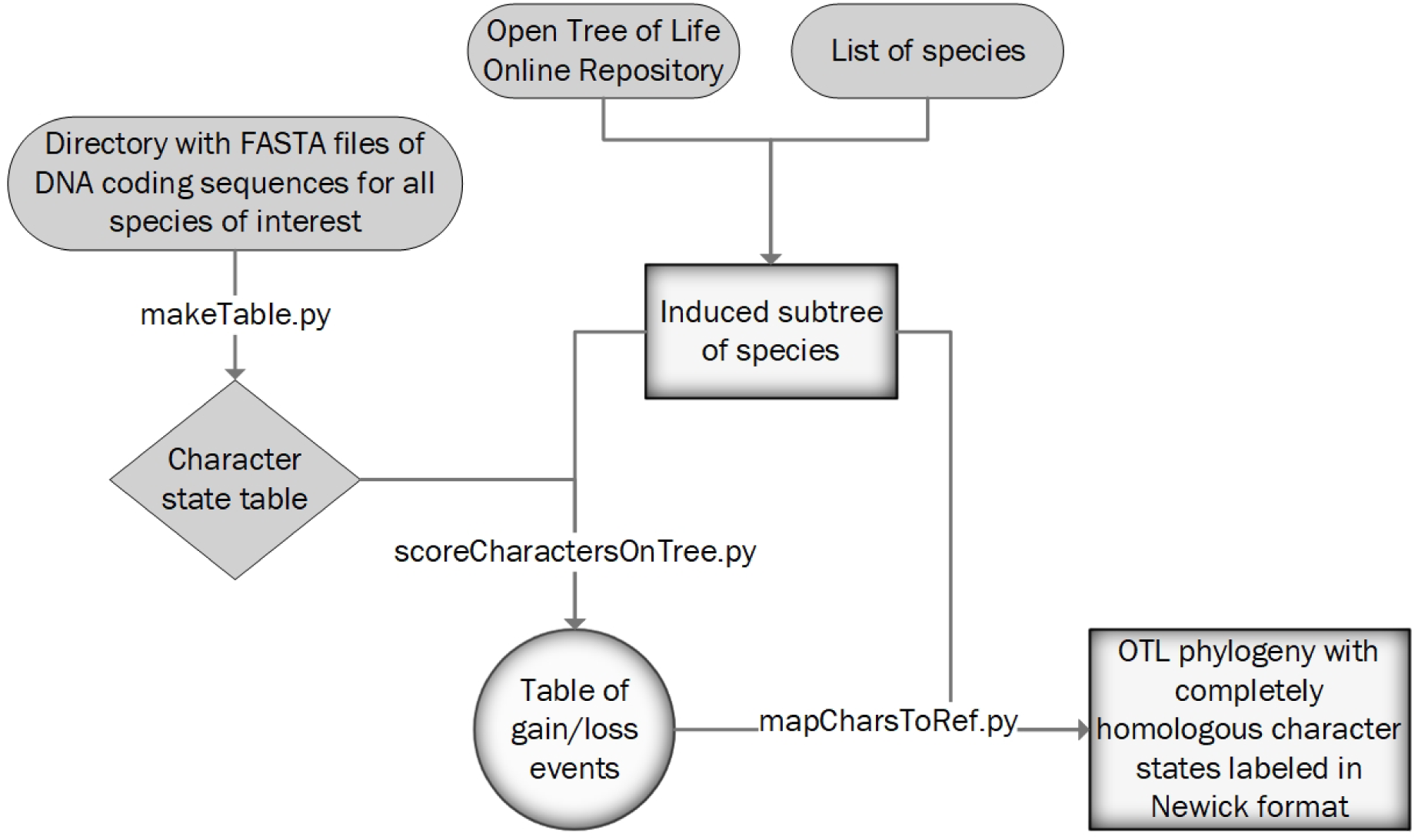
Process for mapping completely congruent character states to the OTL. Starting with a directory where each species has a single FASTA file, a character state table is made, which annotates binary codon usages for all species. This table is passed to an induced subtree from the OTL and creates a table of codon usage transition events. From there, these gains/losses are plotted to the OTL induced subtree, and the phylogeny is reported in Newick format.

### 2.5 Dealing with Limitations in Ortholog Annotations

While the NCBI gene annotations often span hundreds of species, many genes are annotated in fewer than 10 species, with the smaller species partition (i.e., species either using or not using a codon in an ortholog) containing two or three species. These taxonomic distributions were also included in the main analysis. However, the statistical probability (outlined above) of changes in each codon’s usage being congruent with the phylogeny outlined in the OTL for these groups allows for many false positives. For instance, if an ortholog is annotated in four species, with two species using a codon and two species not using that codon, the statistical probability of that codon usage being congruent with the OTL by random chance is 0.33333. Across all species, 9 990 codons fall under this taxonomic distribution, meaning 3 330 of these codons are expected to agree with the OTL by random chance. Although the observed congruence is much higher (4 915), we wanted to ensure that the signal was not simply due to missing ortholog annotations. So, we excluded taxonomic distributions where the probability of obtaining congruence with the OTL was less than one divided by the number of parsimony informative characters. By doing this analysis, we limited the maximum total number of expected congruent codons to one, while ensuring that all observed congruences were statistically unlikely to occur by random chance (i.e., not due to missing data).

### 2.6 Using Codon Aversion to Resolve Controversial Nodes

Since this method uses conditional probability to determine the probability that codon usage follows a proposed phylogeny, we can use this method to compare different phylogenies consisting of the same species. Where two or more contradictory trees exist, each tree is analyzed using the statistical validation described in section 2.2. The resulting t-statistics are then compared, and our method lends support to the phylogeny with the highest t-statistic. We use this process to evaluate the controversial placement of turtles on the OTL.

## 3. Results

### 3.1 Statistical Test

We report the number of different taxonomic distributions, the number of codons with no parallelisms or reversals on the OTL, the *t*-statistic for each group of species used in our analysis, and the *p*-value in Table 1. All taxonomic distributions, expected values, and observed values for each group of species are found in Supplementary Tables 10-18. All 64 codons had similar proportions in the group of codons that mapped to a single gain/loss event on the OTL (*t*-statistic=0.17907, *p-*value=1.0). The ratio of each codon to the total number of codons with a single gain/loss event is depicted in Figure 3.

**Figure 3:**
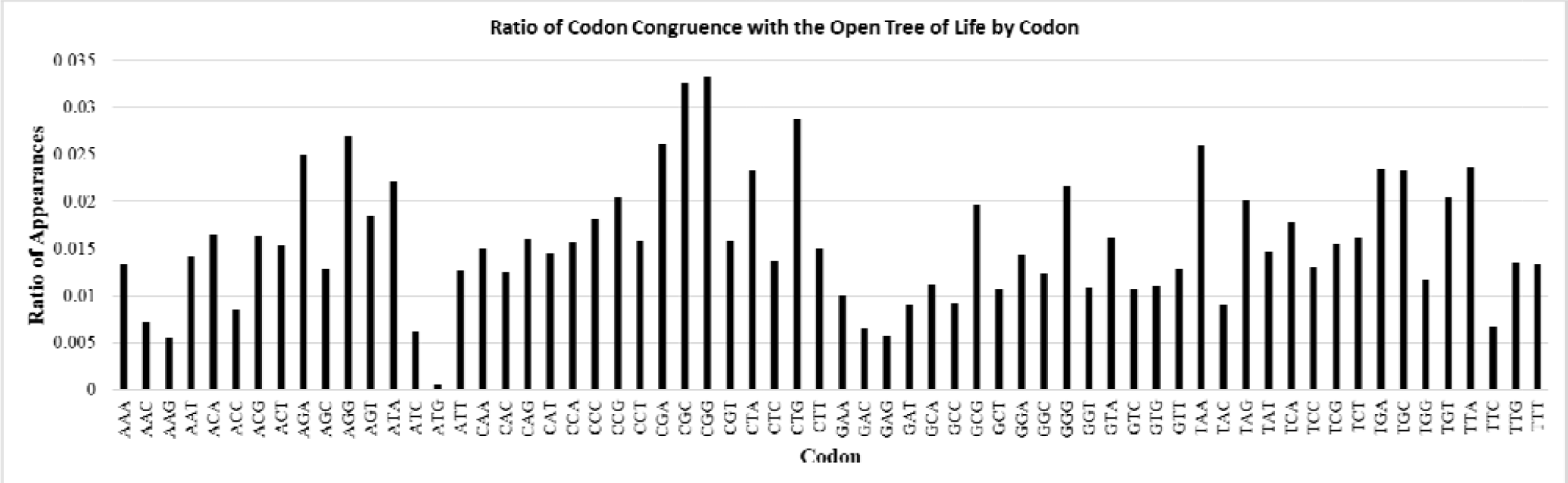
The ratio that each codon with a usage congruent to the OTL. If all codons were given equal weight, the null ratio would be 1.0 / 64 = 0.015625. Observed ratios do not statistically vary from the null, meaning that if a codon usage is congruent with the species relationships outlined in the OTL, it is equally likely for it to be any of the codons.

### 3.2 Permutations

Random permutations for each taxonomic group show that codon usages in archaea, bacteria, plants, mammals, and other vertebrates are not likely to be congruent with the OTL by random chance. The *t-statistics* for the codons with no parallelisms or reversals within these taxonomic groups proposed by the OTL were orders of magnitude larger than the highest *t-statistic* obtained by the random permutations. The largest difference occurred in other vertebrates, where the observed *t-statistic* was 1.50814 × 10^25^ and the highest *t-statistic* from random permutations was 1.16996 × 10^3^.

Although most taxonomic groups had observed *t-statistics* for codon usage that were much greater than those *t-statistics* calculated from random permutations, fungi, invertebrates, and protozoa did not. The *t-statistic* obtained for protozoa was within one order of magnitude of the *t-statistic* of the most improbable random permutation. For fungi, 16% of random permutations had *t-statistics* greater than or equal to observed *t-statistics* from mapping codons to the OTL. Permutations for invertebrates produced *t-statistics* greater than or equal to mapping codons to the OTL 3% of the time.

### 3.3 Missing Ortholog Annotations

Table 2 shows the number of codons within each taxonomic group that have ortholog annotations spanning many species and are unlikely to be completely congruent with the OTL by random chance. Using the statistical validation outlined above, we set a cutoff of one divided by the total number of parsimony informative codons. This ensured that if all codons had a probability less than or equal to the cutoff, at most one codon will be completely congruent with the OTL by random chance. However, as shown in column 4 of Table 2, the maximum number of codons expected to be congruent with the OTL assuming each codon had the maximum probability (i.e., column 3 divided by column 2) was always less than one. By dividing the observed number of codons that agree with the OTL (column 5) by the maximum expected number of codons agreeing with the OTL (column 4), we see a substantial difference in the observed versus the expected in most taxonomic groups (column 5). No orthologs spanned a sufficient number of species in fungi or protozoa to make this analysis possible for those taxonomic groups. Furthermore, few orthologs were annotated across a sufficient number of invertebrates to assess the quality of codon homology in that taxonomic group. The observed number of codons congruent with the OTL in orthologs spanning many species of archaea, bacteria, mammals, other vertebrates, all species, or plants were 37×, 42×, 641×, 985×, 1 109×, and 1 795× larger than the expected values, respectively (column 6).

**Table 2:** Phylogenetic Signal in Orthologs Spanning Many Species. The first column is the taxonomic group analyzed. The second column shows the maximum probability of a codon being completely congruent with the OTL by random chance based on one divided by the total number of parsimony informative characters within that taxonomic group. The third column shows the number of codon characters with a probability less than or equal to the second column. The fourth column shows the maximum number of codons expected to agree with the OTL, assuming all codons in column three had the maximum probability shown in the second column. The fourth column is the product of columns two and three. The fifth column is the number of observed codons that agree with the OTL and have a probability of being congruent with the OTL less than or equal to the maximum probability from the second column. The sixth column is the quotient of the fifth and fourth columns, showing the magnitude difference between the observed and expected codon congruence with the OTL.

| Taxonomic Group | Maximum Probability | Number of Codons with Probabilities Less Than Maximum | Maximum Number of Codons Expected to Agree with the OTL | Number of Codons that Agree with the OTL | Number Observed Divided by Maximum Expected |
| --- | --- | --- | --- | --- | --- |
| All | $1.47453 \times 10^{-6}$ | 470 784 | 0.69419 | 770 | 1 109 |
| Archaea | $6.97058 \times 10^{-5}$ | 8 108 | 0.56517 | 21 | 37 |
| Bacteria | $7.29309 \times 10^{-6}$ | 93 740 | 0.68365 | 29 | 42 |
| Fungi | $1.02155 \times 10^{-5}$ | 0 | 0 | 0 | 0 |
| Invertebrates | $1.07875 \times 10^{-4}$ | 512 | 0.055232 | 0 | 0 |
| Plants | $1.63371 \times 10^{-5}$ | 1 944 | 0.031759 | 57 | 1 795 |
| Protozoa | $1.78508 \times 10^{-5}$ | 0 | 0 | 0 | 0 |
| Mammals | $2.64704 \times 10^{-6}$ | 247 714 | 0.65571 | 420 | 641 |
| Other Vertebrates | $2.83905 \times 10^{-6}$ | 218 129 | 0.61928 | 610 | 985 |

### 3.4 Character States that are Completely Congruent with the OTL

We report the Newick formatted phylogeny from the OTL with all codons that have a singular gain/loss mapped to the trees for each set of species (Supplementary Files 1-9), with the respective character state files showing which codons were gained or lost (Supplementary Files 10-18). Visualizations of the codons that are completely congruent with fungi, invertebrates, plants, protozoa, mammals, and other vertebrates are shown in Supplementary Figures 1-6.

### 3.5 Very Unlikely Character State Distributions

Of the 890 814 codon states analyzed, 25 771 codons had no homoplasy when mapped to the OTL (see Table 1). We further explored a fraction of these character state changes by choosing a subset of codon state changes with a *p*-value ≤ 1 × 10^−25^ of being congruent with the OTL by random chance. Using this arbitrary threshold of 1 × 10^−25^, 52 854 codon characters had taxonomic distributions with a *p*-value ≤ 1 × 10^−25^. Of those characters, the usages of 12 codons were completely congruent with species relationships in the OTL. In Table 3 we report the 12 codons with a p-value ≤ 1 × 10^−25^, and for each codon we report the probability of the taxonomic distribution, the name of the ortholog, and a short description of the species division.

**Table 3:**
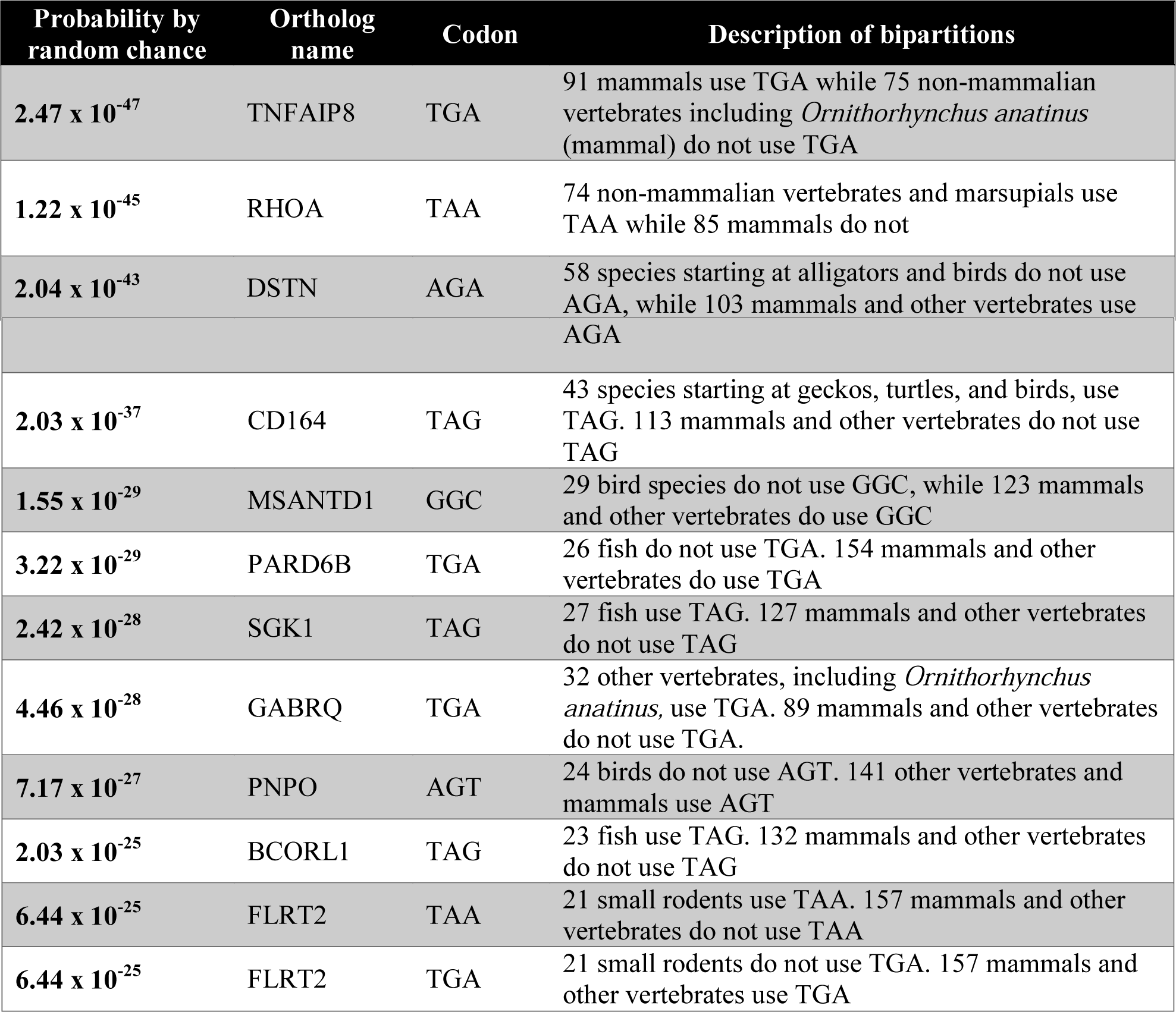
Taxonomic Distributions with a p-value ≤ 1×10-25. The first column is the probability of the taxonomic distribution randomly separating the species according to the OTL classifications. The second column is the name of the orthologous gene. The third column is the codon. The fourth column is a short explanation of how the species were separated.

## 4. Discussion

Using the species relationships reported in the OTL, we identified codons that, once lost or gained, continued in the same character state to all leaf nodes. Two examples of stop codons that persist through evolutionary time from deep nodes to shallow nodes are in the TNFAIP8 (Tumor Necrosis Factor) and RHOA (encodes small GTPase) genes. Both genes play a role in tumor progression, and the specific stop codons used separate most mammals from other vertebrates. Other codons with a singular gain/loss that occurs in deep nodes are outlined in Table 3.

Since each gain/loss event outlined in Table 3 has a probability of occurring by random chance that is less than 1 × 10^−25^ and we studied only 5.2854 × 10^4^ codons that could be congruent with the OTL at that *p*-value threshold, if codon congruence with the OTL were due to random chance, it would be highly unlikely to identify any groups congruent with the OTL. We identified 12 codon usages that were congruent with taxonomic groups found in the OTL. In contrast to the overall analysis where all codons were equally likely to be included as completely congruent with the OTL, in these deep nodal comparisons, nine of the reported codons are stop codons. Since nonsense or nonstop mutations often affect gene function and the stop codon usage persists through time, it is not unreasonable to expect these orthologs to be crucial for species fitness. These codons also lend support to deep species relationships for which these codons map. In conjunction with other methods, codon usage can add support to proposed species trees.

For instance, several controversial nodes were recently analyzed by Shen et al. (2017). In their analysis, they concluded that turtles are the sister taxa to archosaurs (birds and crocodiles) instead of being sister taxa to only crocodiles, and the OTL was updated to reflect this taxonomic relationship. We evaluated this change to the OTL by determining the probability of each tree (with turtles as sister taxa to archosaurs and with turtles as sister taxa to crocodiles) based on codon usages. We used the “other vertebrates” taxonomic group and changed only the location of turtles on the tree. We found that the probability of observed codon usages randomly mapping to the “other vertebrates” tree with turtles as sister taxa to crocodiles was 7.3306 × 10^16^ higher than the probability of turtles being sister taxa to archosaurs. This analysis lends support to keeping turtles as sister taxa to archosaurs on the OTL because the probability of the codons randomly mapping to that tree is smaller than the probability of the codons randomly mapping to the other tree.

We recognize a bias toward recovering shallow nodes using this method because many orthologs are not yet annotated for all species. To overcome this bias, we looked at codon usages that were statistically unlikely to be congruent with the OTL and found that across all species, codons were 1 109x more likely to be congruent with the OTL than expected by random chance. Of the 890 814 parsimony informative codons, 590 366 (66.27%) had ortholog annotations for at least 100 species, and 6 688 (25.95%) of the 25 771 codons that were congruent with the OTL taxonomic relationships were from orthologs annotated in at least 100 species. Furthermore, 11 codons whose usage was congruent with the OTL were identified from orthologous genes that were annotated in more than 1 000 species. Identifying complete codon congruence with the OTL in thousands of groups of at least 100 species shows that homology in codon usage can exist in larger taxonomic groups. By performing random permutations of our dataset, we also show that there is less congruence between the codons that were not congruent with the species relationships in the OTL than with the original codon dataset. This analysis shows that although the majority of codon usages do not have a singular gain/loss when mapped to the OTL, codon usages are more likely to follow species relationships in the OTL than in a random phylogeny.

Although we do not have sufficient ortholog annotations to conclude codon congruence with the OTL in fungi, invertebrates, or protozoa, this analysis shows that codon usage is maintained in archaea, bacteria, plants, mammals, and other vertebrates. We also propose that the framework that we provide for performing this analysis will reveal a phylogenetic signal in the other taxonomic groups when more orthologs are annotated across those species. Looking forward, we anticipate that codon usage will become another tool for evaluating different species trees, similar to our evaluation of the placement of turtles on the OTL.

## 5. Acknowledgements

We appreciate the contributions of Brigham Young University and the Fulton Supercomputing Laboratory at Brigham Young University for supporting our research.

## 6. Competing Interests

The authors declare no competing interests.

## Notes

https://github.com/ridgelab/codon_congruence

