## Supplementary material for "Codon Use and Aversion is Largely Phylogenetically Conserved Across the Tree of Life": Supplementary Information.pdf

**Supplementary Information**
**for**
**Codon Use and Aversion is Largely Phylogenetically Conserved Across the Tree of Life**

**Figures:**

Supplementary Figure 1: Fungi: The Open Tree of Life with annotated character state changes.
Supplementary Figure 2: Invertebrates: The Open Tree of Life with annotated character state changes.
Supplementary Figure 3: Plants: The Open Tree of Life with annotated character state changes.
Supplementary Figure 4: Protozoa: The Open Tree of Life with annotated character state changes.
Supplementary Figure 5: Mammals: The Open Tree of Life with annotated character state changes.
Supplementary Figure 6: Other Vertebrates: The Open Tree of Life with annotated character state changes.

**Tables**

Supplementary Table 1: All species: For each parsimony informative codon character state, the name of the ortholog and codon, the number of gains, the number of losses, the number of unknown gains/losses from the root node, the number of species in the smaller group, the number of total species with that ortholog, the percent of species in the smaller group, and the total number of gains/losses divided by number of species in the smaller group.
Supplementary Table 2: Archaea: See Supplementary Table 1
Supplementary Table 3: Bacteria: See Supplementary Table 1
Supplementary Table 4: Fungi: See Supplementary Table 1
Supplementary Table 5: Invertebrates: See Supplementary Table 1
Supplementary Table 6: Plants: See Supplementary Table 1
Supplementary Table 7: Protozoa: See Supplementary Table 1
Supplementary Table 8: Mammals: See Supplementary Table 1
Supplementary Table 9: Other Vertebrates: See Supplementary Table 1
Supplementary Table 10: All species: The number of species in the smaller clade, the total number of species, the number of clades with this cladal distribution, the number of groups with this cladal distribution which are expected to be consistent with the OTL based on random chance, the number of observed groups that were consistent with the OTL.
Supplementary Table 11: Archaea: See Supplementary Table 10
Supplementary Table 12: Bacteria: See Supplementary Table 10
Supplementary Table 13: Fungi: See Supplementary Table 10
Supplementary Table 14: Invertebrates: See Supplementary Table 10
Supplementary Table 15: Plants: See Supplementary Table 10
Supplementary Table 16: Protozoa: See Supplementary Table 10
Supplementary Table 17: Mammals: See Supplementary Table 10
Supplementary Table 18: Other Vertebrates: See Supplementary Table 10

**Files**

Supplementary File 1: All Species: Species relationships from the OTL annotated in Newick format with homologous character state changes labeled. A range is given, which points to the order in the accompanying character state change file.
Supplementary File 2: Archaea: See description for Supplementary File 1
Supplementary File 3: Bacteria: See description for Supplementary File 1
Supplementary File 4: Fungi: See description for Supplementary File 1

Supplementary File 5: Invertebrates: See description for Supplementary File 1
Supplementary File 6: Plants: See description for Supplementary File 1
Supplementary File 7: Protozoa: See description for Supplementary File 1
Supplementary File 8: Mammals: See description for Supplementary File 1
Supplementary File 9: Other Vertebrates: See description for Supplementary File 1
Supplementary File 10: All Species: Accompanying character state change file for Supplementary File 1.
Supplementary File 11: Archaea: Accompanying character state change file for Supplementary File 2.
Supplementary File 12: Bacteria: Accompanying character state change file for Supplementary File 3.
Supplementary File 13: Fungi: Accompanying character state change file for Supplementary File 4.
Supplementary File 14: Invertebrates: Accompanying character state change file for Supplementary File 5.
Supplementary File 15: Plants: Accompanying character state change file for Supplementary File 6.
Supplementary File 16: Protozoa: Accompanying character state change file for Supplementary File 7.
Supplementary File 17: Mammals: Accompanying character state change file for Supplementary File 8.
Supplementary File 18: Other Vertebrates: Accompanying character state change file for Supplementary File 9.
