## Supplementary figures and images for "Codon Use and Aversion is Largely Phylogenetically Conserved Across the Tree of Life"

### Supplementary Figure 1 labeled fungi.pdf

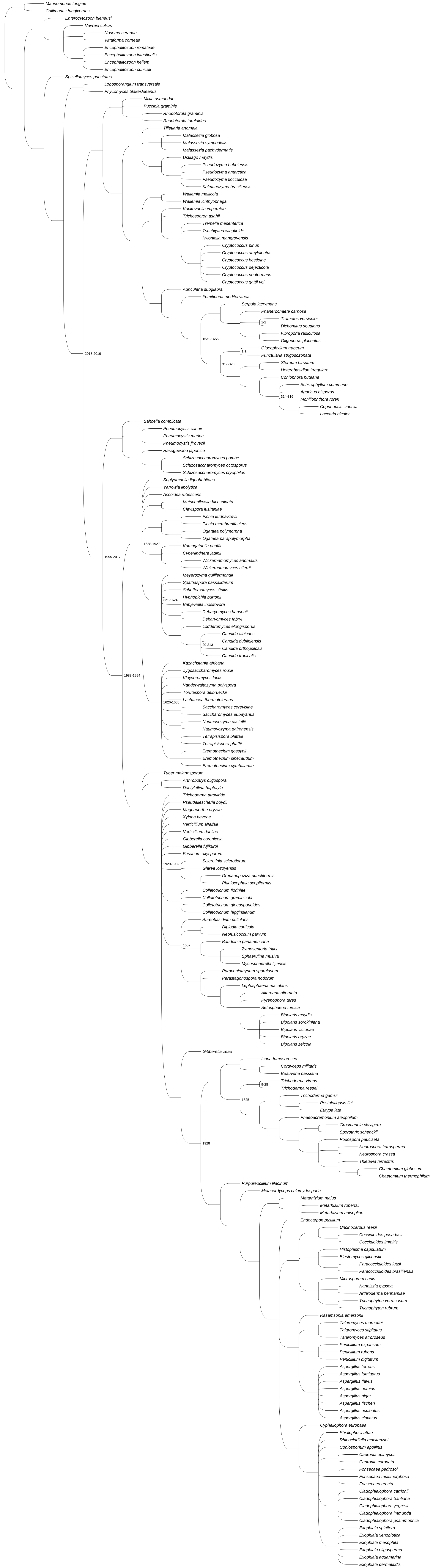

### Supplementary Figure 2 labeled invertebrates.pdf

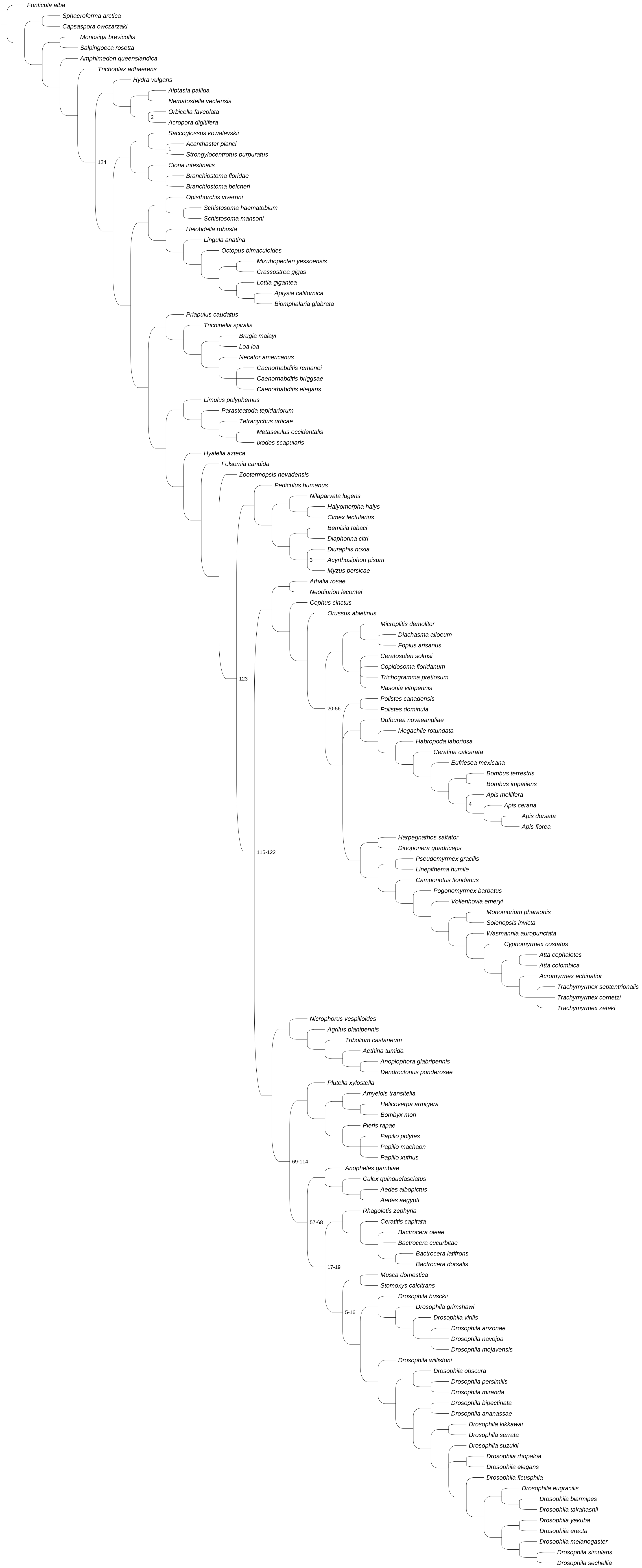

### Supplementary Figure 3 labeled plants.pdf

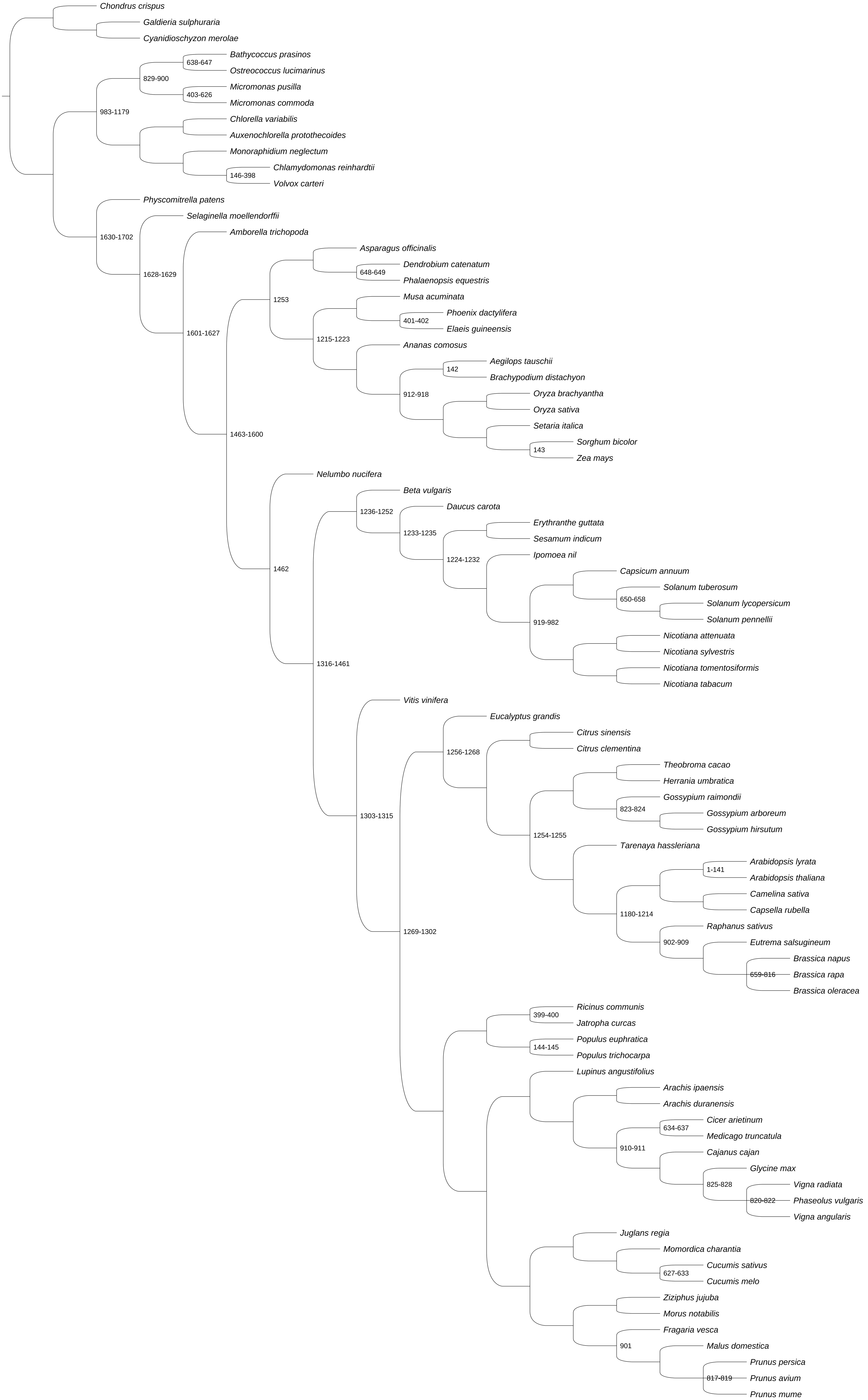

### Supplementary Figure 4 labeled protozoa.pdf

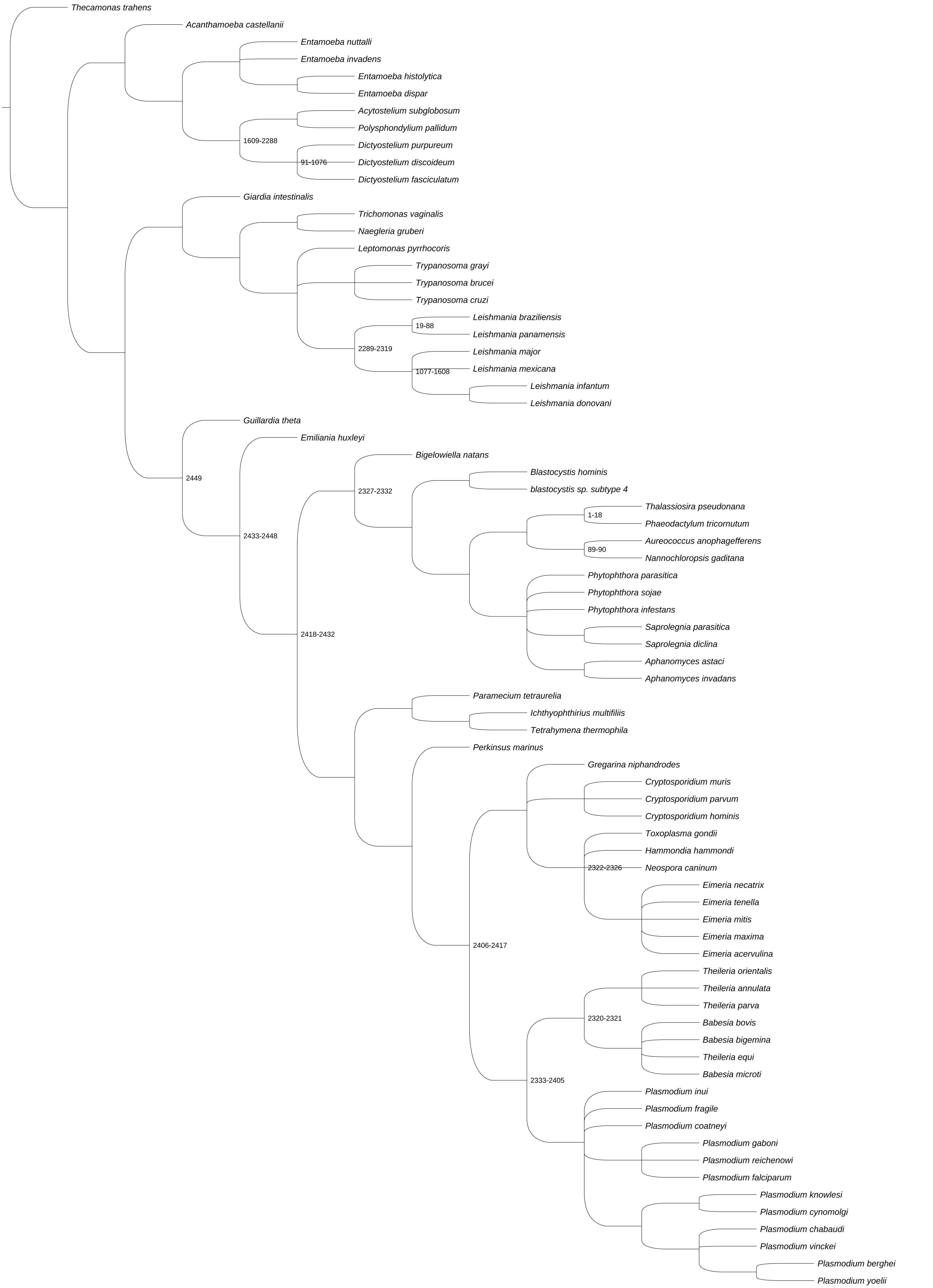

### Supplementary Figure 5 labeled mammals.pdf

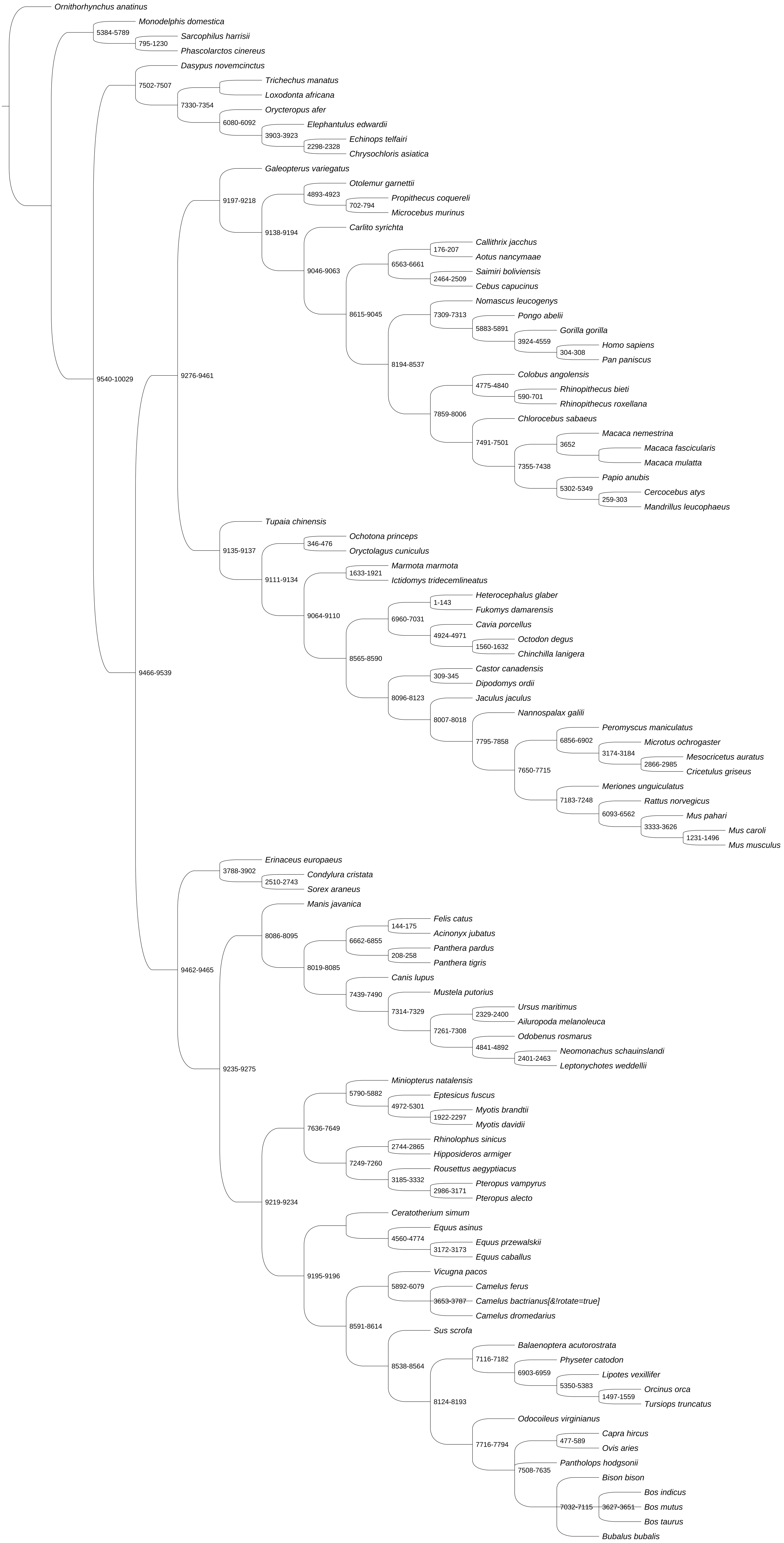

### Supplementary Figure 6 labeled other vertebrates.pdf

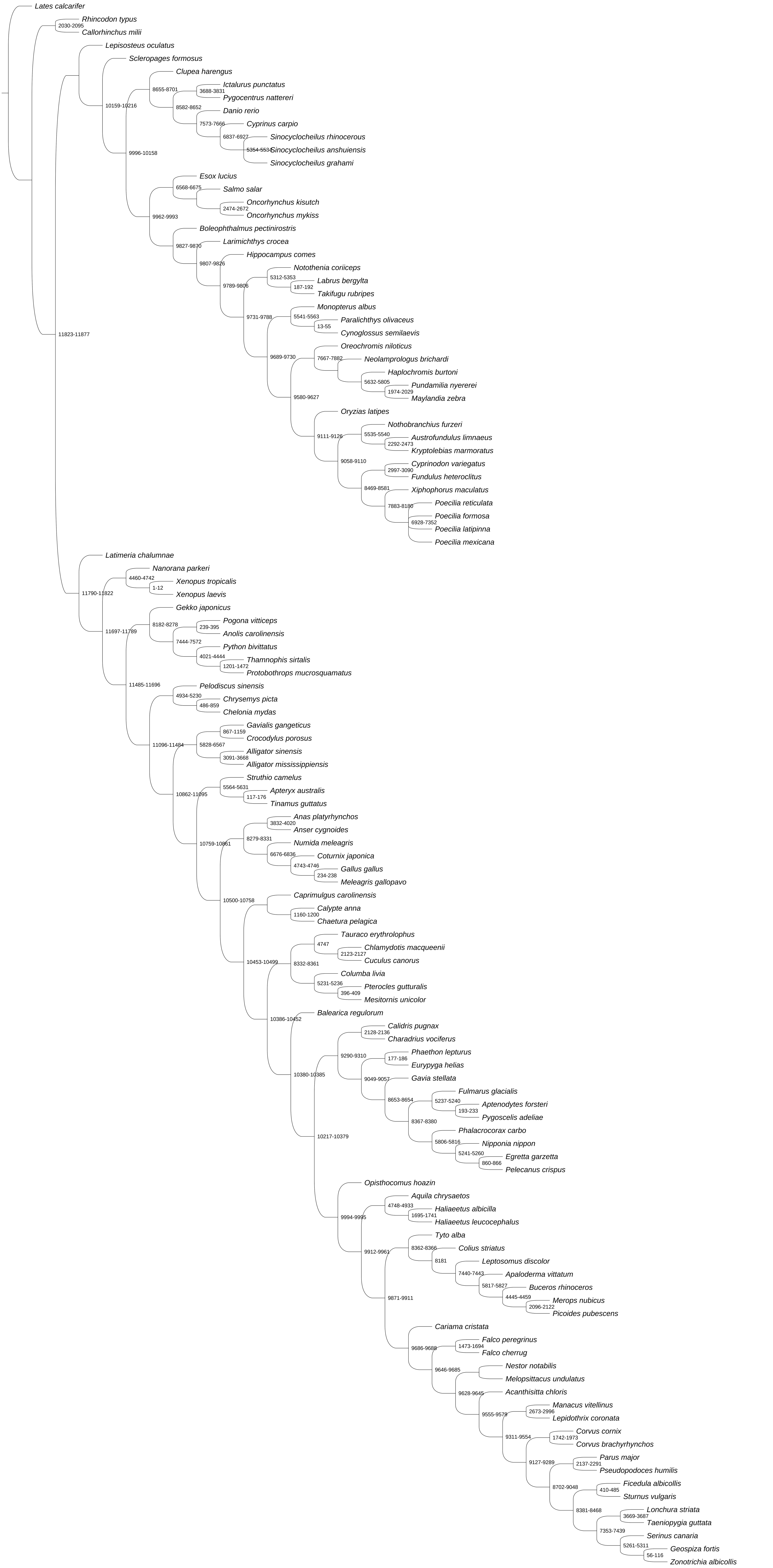
